# Spatiotemporal Dynamics of Anionic Phospholipids Orchestrate Lateral Root Initiation and Morphogenesis in *Arabidopsis thaliana*

**DOI:** 10.1101/2025.08.08.669267

**Authors:** Joseph G. Dubrovsky, Juan Li, Sami Bouziri, Eric Bormann, Celine Geiger, Jazmin Reyes-Hernandez, Alexis Maizel

**Affiliations:** Center for Organismal Studies (COS), University of Heidelberg, Heidelberg, Germany; Laboratory of Developmental Biology of Plant Roots at the Instituto de Biotecnología, Universidad Nacional Autónoma de México (UNAM), Cuernavaca, Mexico; Guizhou University of Traditional Chinese Medicine, School of Basic Medicine, Guizhou, PRC – [JRH] Department of Plant and Environmental Sciences, University of Copenhagen, Copenhagen, Denmark

## Abstract

Lateral root (LR) development in *Arabidopsis thaliana* requires precise coordination of pericycle founder cell (FC) specification, patterning, and morphogenesis. While auxin signalling is well established in this process, the role of membrane lipid signalling—particularly phosphoinositides—remains less understood. Here, we investigate the contribution of the anionic phospholipids PI4P, PI(4,5)P₂, and phosphatidylserine (PS) to LR formation using live-cell biosensors, genetic mutants, and inducible lipid depletion tools. We show that PI4P is uniformly distributed throughout lateral root primordia (LRPs), whereas PI(4,5)P₂ is specifically depleted in the proliferative core during early LRP development. Time-lapse imaging revealed stable PI4P and PI(4,5)P₂ levels before and after FC activation, while PS increased rapidly post-activation. In xylem-pole pericycle (XPP) cells, PI(4,5)P₂ decreased and PS increased following LR initiation, with both changes occurring in a membrane-domain-specific manner. Genetic analysis of the *pip5k1pip5k2* double mutant, deficient in PI(4,5)P₂ synthesis, revealed impaired LR initiation and emergence. Conversely, inducible depletion of PI(4,5)P₂ using the iDePP system enhanced LRP initiation and accelerated development when activated after FC specification. These results suggest that PI4P functions as a stable basal lipid, while PI(4,5)P₂ and PS undergo dynamic, spatially regulated changes critical for LR progression. Notably, PI(4,5)P₂ acts as a negative regulator of LRP initiation and morphogenesis. Our findings highlight how lipid signalling, in coordination with hormonal cues, provides spatial and temporal control over pericycle cell behaviour and lateral root organogenesis.

**SIGNIFICANCE STATEMENT:** This study reveals how specific membrane lipids help control where and when new lateral roots form in plants. While plant hormones like auxin are known to guide root branching, this work shows that lipids such as PI(4,5)P₂ and phosphatidylserine also play key roles. Using live imaging and genetic tools, we found that reducing PI(4,5)P₂ in certain root cells promotes root growth by triggering cell division and development. In contrast, phosphatidylserine level increases right when root-forming cells become active. These discoveries highlight a new layer of control in plant development and suggest that lipids help fine-tune the formation of roots, which is essential for how plants take up water and nutrients.

## INTRODUCTION

The pericycle is a unique plant tissue that retains proliferative capacity outside the root apical meristem, contributing to both lateral root (LR) formation and secondary growth. Through its ability to give rise to new meristems, including vascular cambium and phellogen, the pericycle plays a central role in root system architecture and radial expansion (1–3). Despite its developmental importance, the cellular physiology of pericycle cells—and in particular the molecular mechanisms that govern their participation in LR initiation—remains incompletely understood.

Analysis of proliferation activity of pericycle cells outside the root apical meristem (RAM) showed that when specific xylem pole pericycle (XPP) cells acquire LR founder cell (FC) identity. These cells start LR primordium (LRP) formation, no quiescence is found, as pericycle cells continue cycling. The average cell cycle time in the RAM for XPP cells is the same as the period from their departure from the RAM till the early LRP stage (1). Importantly, while some XPP cells become LR FCs, other XPP cells simply divide in the root differentiation zone, become shorter, and do not enter the LR formation program (1, 2). In *A. thaliana,* LR initiation events take place in an acropetal sequence, meaning that a new LRP is initiated towards the root apex in relation to a previously initiated LRP (4). This suggests that the physiological state of XPP cells before and after LRP initiation is different. In addition, anticlinal divisions of XPP cells that do not participate in LRP formation are followed by periclinal divisions, which impact vascular and cork cambium formation (3, 5).

Anionic phospholipids, including phosphatidic acid, phosphatidylserine (PS), and phosphoinositides, comprise a minor fraction of plasma membrane phospholipids but have critical roles in regulating membrane properties and signalling. These lipids contribute to membrane identity by altering surface charge and curvature, and they act as dynamic regulators of protein localisation and activity (6–8). In plants, anionic lipids are increasingly recognised as key modulators of developmental processes, including cell polarity, division, vesicle trafficking, and gene expression (6, 9). Among these lipids, PS, phosphatidylinositol 4-phosphate (PI4P) and its derivative phosphatidylinositol 4,5-bisphosphate [PI(4,5)P₂] are of particular interest for their role in development. They are essential for maintaining membrane electrostatics (10).

PS-specific distribution between endosome and plasma membrane depends on cell differentiation state, and in the elongation zone, PS becomes more abundant in endosomes than at the plasma membrane (10). However, in the presence of auxin, PS becomes predominantly localised in the plasma membrane in epidermal cells of the young elongation zone (11). As LR FCs are known to accumulate auxin (12), one can speculate that PS abundance at the plasma membrane of FCs may change after FC specification.

The synthesis of PI(4,5)P₂ from PI4P is catalysed by phosphatidylinositol monophosphate 5-kinases (PIP5Ks), of which *Arabidopsis thaliana* possesses eleven isoforms (13). Several of these are expressed in roots and have been implicated in diverse aspects of root development. For example, PIP5K1 and PIP5K2 regulate cell elongation, vascular differentiation, and root growth (14–16), while PIP5K2 also influences LR formation and gravitropic response via modulation of the auxin transporter PIN2 (17). Importantly, PIP5K2 is expressed in LRPs, and the application of PI(4,5)P₂ can rescue LR defects in *pip5k2* mutants—yet the precise role of PI(4,5)P₂ in LR development remains unknown.

In addition to synthesis, PI(4,5)P₂ turnover is also crucial. The PI(4,5)P₂ phosphatase SAC9 restricts PI(4,5)P₂ accumulation during cytokinesis, and its loss results in root growth defects due to impaired cell proliferation (18, 19). These findings point to the importance of balanced PI4P and PI(4,5)P₂ levels in root development. However, despite these advances, no prior studies have systematically examined the distribution or functional role of anionic phospholipids in the pericycle during LR initiation and morphogenesis. LRP morphogenesis is a relatively stereotypic process that starts from anticlinal divisions of FCs followed by periclinal divisions, giving rise to well-described LRP developmental stages (20) whose development is under a complex hormonal and genetic control (21). LR morphogenesis is accompanied by different cell proliferation potential of core and flanking LRP cells, resulting in the formation of a typical dome-shaped LRP (22, 23).

Here, we investigated the spatial dynamics and functional importance of anionic phospholipids—particularly PI4P, PI(4,5)P₂, and PS—during the early stages of LR development. Using live-cell lipid biosensors, loss-of-function mutants, and inducible depletion systems, we revealed that pericycle cells exhibited distinct phospholipid distributions before and after LR initiation and during FC activation. We demonstrated that PI(4,5)P₂ synthesis is essential not only for LRP initiation but also for LR emergence, and that localised depletion of PI(4,5)P₂ in founder-cell domains promotes LRP formation. Our findings highlight a critical role for spatial PI(4,5)P₂ distribution in coordinating pericycle participation in LRP formation and LR organogenesis. Our work revealed that this and other anionic phospholipids are essential players in root system formation.

## RESULTS

### Distinct PI(4,5)P**₂** Distribution Patterns Reveal Local Regulation During Lateral Root Primordium Development

In *A. thaliana* root, the distribution of PI4P and PI(4,5)P₂ was studied mainly in the epidermis, where they are most abundant at the plasma membrane and at a lower level in the Golgi and post-Golgi/endosomal compartments (24). To gain insight into the distribution of PI4P and PI(4,5)P₂ during the initial stages of LRP formation, we monitored their presence and abundance using sensor lines specific to each lipid type, which consist of CITRINE-tagged lipid-binding domains (LBDs). We used the ubiquitously expressed reporter *UBQ10prom::CITRINE-1xPH^OSBP^*for PI4P and *UBQ10prom::CITRINE-2xPH^PLC^* for PI(4,5)P₂ (24).

We measured the plasma membrane and cytoplasmic signals in the LRP core, flanking cells, and adjacent pericycle cells. Due to the known exclusion of PI(4,5)P₂ from the cell plate during division (19, 25), we restricted membrane measurements to the periclinal sides facing the endodermis (outer) and protoxylem (inner). All signals were normalised to the LRP core region. As both lipids accumulated at the plasma membrane and intracellularly (thereafter cytosolic signal), we quantified both pools.

While PI4P was evenly distributed across all LRP cells, from Stage I to IV, with no difference between the core and flanking domains (Fig. 1A, C), PI(4,5)P₂ exhibited a dynamic pattern. Initially equally abundant in the central and flanking regions of Stage I primordium, PI(4,5)P₂ abundance at the membrane decreased in the primordium core by comparison to its flanks and adjacent pericycle cells from Stage II onwards (Fig. 1B, D). A similar trend was found when considering the cytoplasmic signal (Figure S1). These findings suggest that PI(4,5)P₂ is downregulated specifically in proliferative core cells of the LRP. Additionally, its elevated levels in adjacent pericycle cells imply that PI(4,5)P₂ reduction may be associated with activation of FCs during LRP initiation.

**Figure 1.**
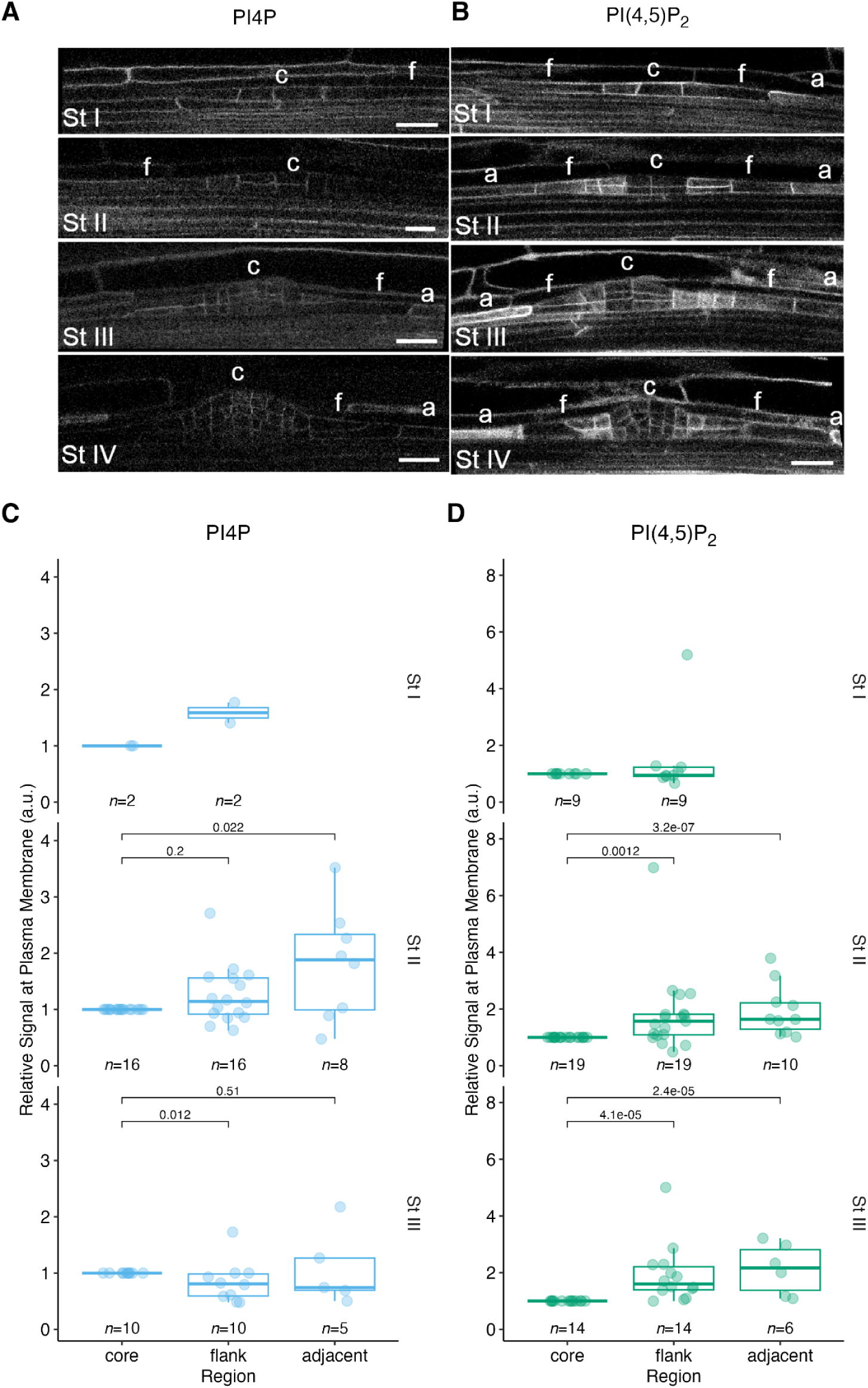
Spatial distribution of PI4P and PI(4,5)P₂ at the plasma membrane during lateral root primordium (LRP) development. (A–B) Confocal images showing plasma membrane and cytoplasmic localisation of biosensors for PI4P *UBQ10prom::CITRINE-1xPHOSBP* (A) and for PI(4,5)P₂ *UBQ10prom::CITRINE-2xPHPLC* (B) biosensors in LRPs at developmental stages I–IV. Signal distribution is shown across the LRP core (c), flanking (f), and adjacent (a) pericycle cells. Scale bars = 10 µm. (C–D) Quantification of biosensor signal at the plasma membrane in the LRP core, flanks, and adjacent pericycle cells at developmental stages I to III. Signal intensity is expressed as relative fluorescence (a.u.), normalised to the core. (C) PI4P signal remains relatively uniform across core, and adjacent regions, while slightly reduced at stage III between core and adjacent cells. (D) PI(4,5)P₂ signal is significantly reduced in the LRP core compared to flanking and adjacent pericycle cells at both stages II and III, indicating spatial depletion of PI(4,5)P₂ within the proliferative core region. Each point represents a single cell; *n* indicates the number of cells analysed. Boxplots show medians and interquartile ranges. *P*-values from a non-parametric Kruskal-Wallis one-way analysis of variance on ranks. See also Figure S1.

### PI(4,25)P**₂** and PS Display Opposing Plasma Membrane Dynamics in XPP Cells After LRP Initiation

Since LRP initiation in *A. thaliana* takes place in a strictly acropetal pattern (4, 12), we hypothesised that if anionic phospholipids play a role in LR initiation and development, their abundance and/or distribution in XPP cells should change after LR initiation compared to those in XPP cells before LR initiation. To verify this hypothesis, the most distal LRP was detected and the biosensor level of a specific anionic phospholipid was measured in XPP cells before (distally to the most distal LRP) and after LR initiation (Fig. 2A). To achieve cell-type-specific resolution, we generated reporter lines using the XPP-active promoter *pXPP* (26) to drive expression of red fluorescent protein mScarlet fused to lipid-binding domains specific for PI4P and PI(4,5)P₂. Given the role of phosphatidylserine (PS) in regulating plasma membrane organisation, we expanded our analysis to PS. These lines were generated in a background expressing a constitutive plasma membrane marker (*pUBI:3xGFP:PIP1;4, (27)*) for normalisation, and a nuclear marker (*H2B:3xmCherry*) expressed under XPP and FC-specific promoter, *pGATA23* (28) to easily identify XPP cells and their derivatives (Figure 2A). To quantitatively assess the abundance of these phospholipids, we measured the ratio of lipid nanosensor signal (mCherry) to the GFP signal intensity along different membrane domains—specifically, the longitudinal inner (facing the protoxylem or procambium cells) and outer (facing the endodermis) walls, and adjacent to the transverse cell walls of XPP cells.

**Figure 2.**
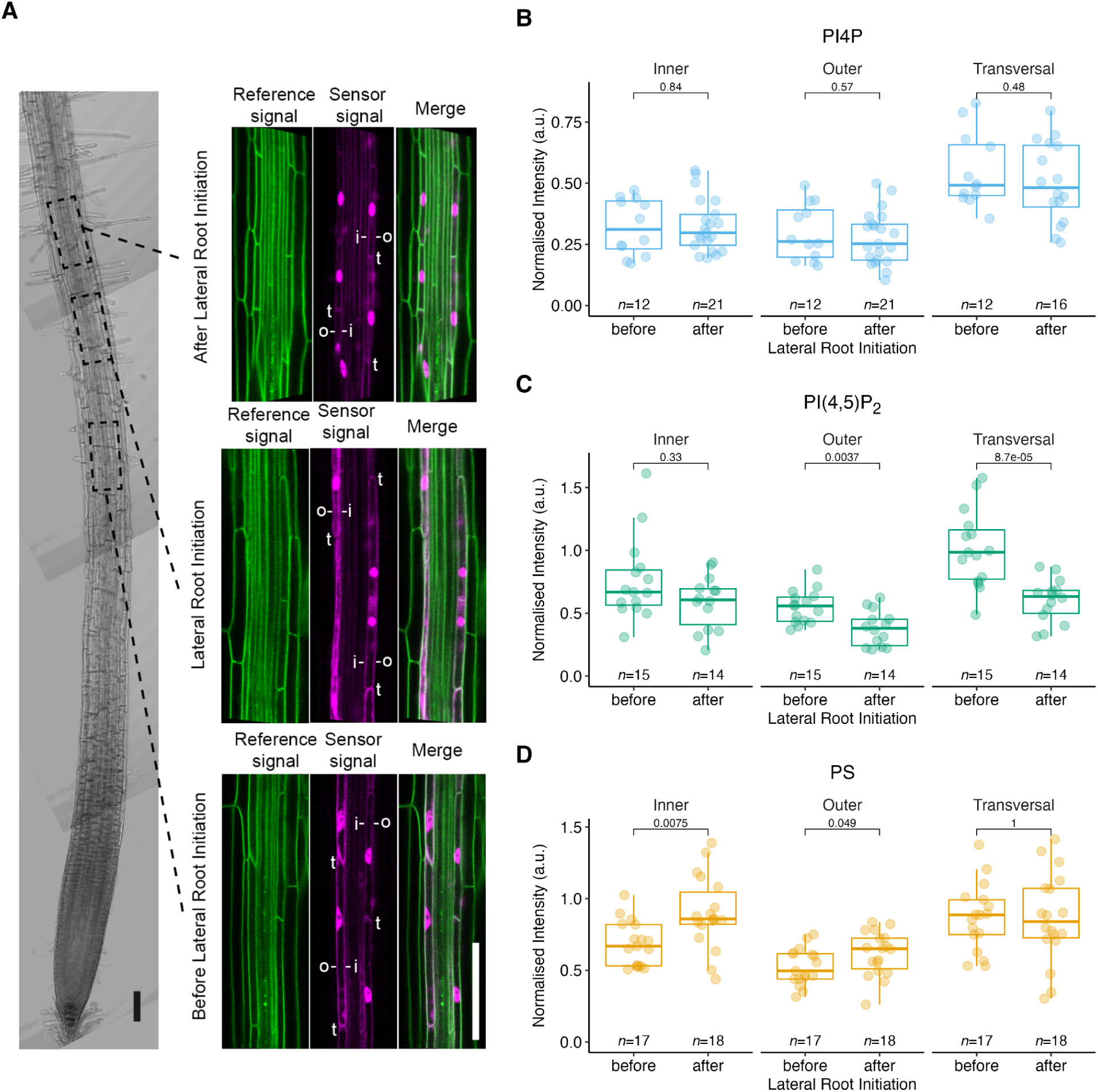
Changes in anionic phospholipid abundance in xylem-pole pericycle cells before and after lateral root initiation. (A) Confocal images showing the localisation of lipid biosensors in xylem-pole pericycle (XPP) cells before, during, and after lateral root initiation. Sensor plasma membrane signal (magenta) is shown alongside the plasma membrane reference signal (green, *pUBI::3xGFP:PIP1;4*), with merged images at right. Nuclear positions (magenta) are marked by *pGATA23::H2B:3xmCherry*. Measurements were taken at the inner (i), outer (o), and transverse (t) sides of XPP cell plasma membranes in defined zones: before LRI (distal to the most distal LRP) and after LRI (just shootward to the most distal LRP). Lateral root initiation shows Stage 0 (the first division of founder cell; (29)). Scale bars = 50 µm. (B–D) Quantification of normalised sensor signal intensity for PI4P (B), PI(4,5)P₂ (C), and phosphatidylserine (PS) (D) before and after LRI at inner, outer, and transversal cell wall domains. (B) PI4P levels remain unchanged across all membrane domains before and after LRI (*P* > 0.4 for all). (C) PI(4,5)P₂ levels significantly decrease after LRI at the outer and transverse membranes (*P* = 0.0037 and *P* < 0.0001, respectively), while inner membrane levels remain unchanged. (D) PS levels significantly increase after LRI at the inner (*P* = 0.0075) and outer (*P* = 0.049) membranes, with no change at the transverse side. Each dot represents an individual membrane measurement; boxplots show median and interquartile ranges. *n* values indicate the number of cells analysed. *P*-values from a non-parametric Kruskal-Wallis one-way analysis of variance on ranks. The lines used for this analysis were: JL012 (for PI4P), SB007, SB008, SB009 (for PI(4,5)P₂), SB043, SB044, SB045 (for PS). See Table S1 for details.

PI4P distribution at the plasma membrane of XPP cells remained unchanged before and after LRP initiation, indicating that PI4P levels are stable during this developmental transition (Figure 2B). In contrast, PI(4,5)P₂ abundance in XPP cells after LR initiation decreased in particular at the outer and transverse domains to approximately 69% (outer domain) and 59% (transverse domain) of their pre-initiation levels (Figure 2C). Interestingly, the distribution pattern of PS differed. Following LRP initiation, PS abundance in the inner domain increased (Figure 2D), but remained the same in the outer and transversal domains. Together, these results reveal that the abundance of each phospholipid responds differently to LRP initiation and their response is domain-specific. While PI4P remains unchanged, PI(4,5)P₂ decreases, and PS increases in a domain-specific manner. These observations suggest that phospholipid composition is dynamically regulated during the early stages of LR development. The specific decrease in PI(4,5)P₂, despite stable PI4P levels, implies a targeted regulatory mechanism that may contribute to LR initiation and LRP formation.

### The timing of PIP4, PI(45)P_2_, and PS modulation is different upon LR founder cell activation

Our analysis of PI4P and PI(4,5)P₂ distribution in developing LRPs (Figure 1) showed that while PI4P levels remained uniform across the core and flanks, PI(4,5)P₂ was significantly reduced in the LRP core. This reduction may be associated with high proliferative activity in core cells during morphogenesis. Similarly, changes in PI(4,5)P₂ and PS levels in XPP cells after LR initiation (Figure 2) suggest that dynamic regulation of these anionic phospholipids may play a role in LR initiation. To examine the dynamics of the changes in the abundance of these lipids, during, and after FC activation, we performed time-lapse imaging using biosensor lines specific for PI4P (JL002: *pGATA23::mScarlet:FAPP1*; JL001: *pXPP::mScarlet:FAPP1*), PI(4,5)P₂ (SB001: *pXPP::mScarlet:2xPH(PLC)*), and PS (SB037: *pXPP::mScarlet:1xC2Lact*; SB038: *pGATA23::mScarlet:1xC2Lact*). These lines were crossed with auxin-responsive reporters *pDR5rev::GFP* (*30*) or *pDR5rev::3xVENUS-N7* (*31*), and F1 plants were analysed in time-lapse experiments. Auxin response, reported by *pDR5*, increases gradually along the root’s central cylinder, reaching a peak at the LR initiation zone near young protoxylem cells (32). In this region, presumptive FCs were identified and imaged every 0.5 to 1.5 hours for 11 to 16 hours. FCs that proceeded to divide and develop into LRPs were analysed for biosensor signal intensity to detect short-term changes in PI4P, PI(4,5)P₂, and PS abundance (Figure 3). Signal intensities were normalised to the value at the time of the first FC division, which marked the point of FC activation.

**Figure 3.**
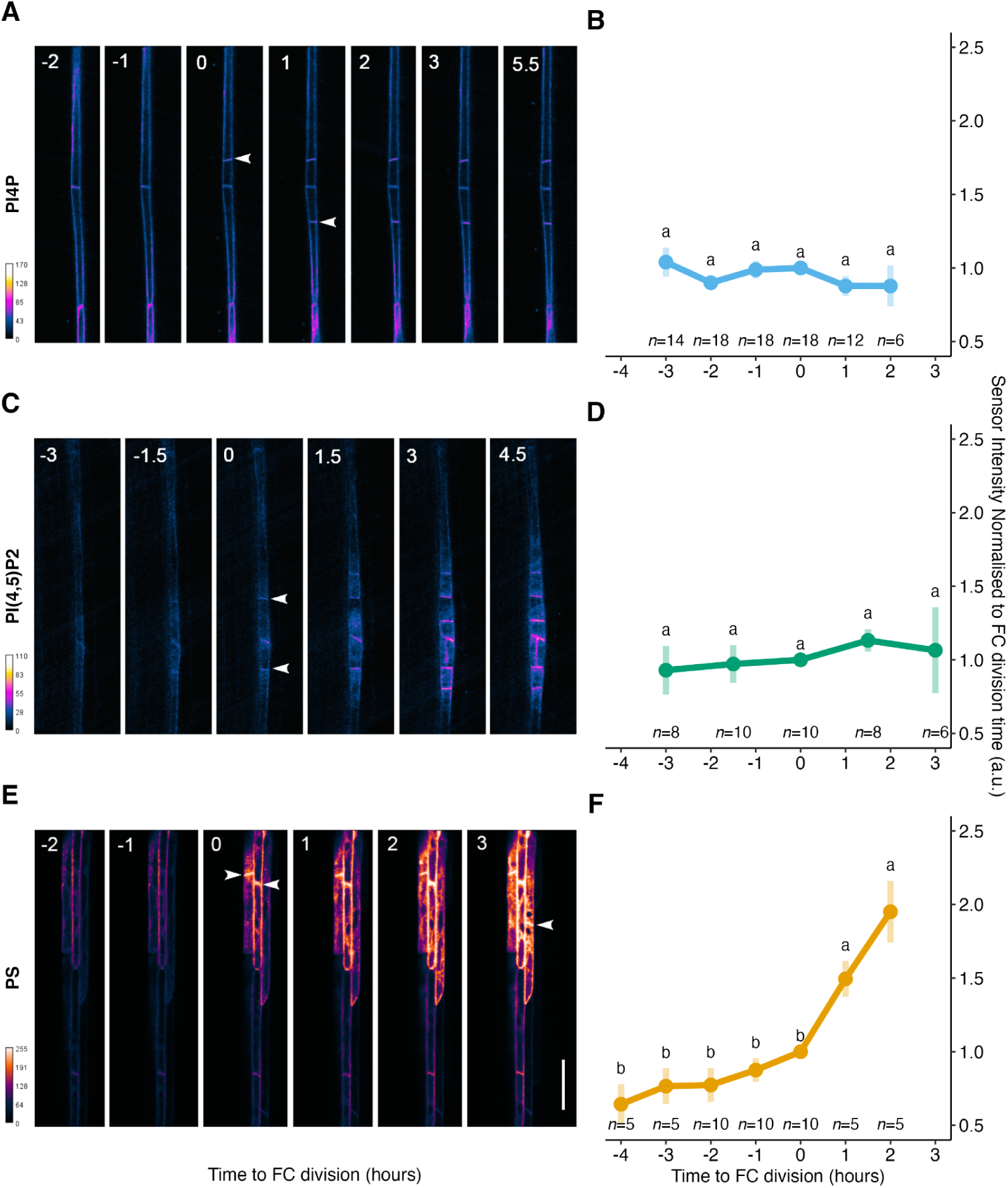
Time-resolved analysis of anionic phospholipid abundance during founder cell (FC) activation. (A, C, E) Representative time-lapse confocal images showing signal dynamics of biosensors for PI4P (A), PI(4,5)P₂ (C), and phosphatidylserine (PS) (E) in xylem-pole pericycle (XPP) cells before and after FC activation. Time is shown in hours relative to the first division of the FC (time 0). Arrowheads mark the dividing FC. Color scales represent relative signal intensity (in arbitrary units, a.u.). Scale bar = 20 µm. (B, D, F) Quantification of biosensor signal intensity over time, normalised to the signal at time 0 (first FC division), for PI4P (B), PI(4,5)P₂ (D), and PS (F). PI4P and PI(4,5)P₂ levels remain stable before and after FC division (B, D), with no significant changes detected. In contrast, PS levels show a marked increase after FC division, with signal intensity nearly doubling by 2–3 hours post-division (F), indicating rapid post-activation enrichment. Each point represents the mean ± SEM; *n* indicates the number of FCs analysed at each time point. Different letters indicate statistically significant differences between time points (*P* < 0.05, ANOVA with post hoc test). Lines used: JL002 x DR5revVENUS (A), SB001 x DR5revGFP (C), and SB037 x DR5revGFP (E). Note, only the biosensor red channel is shown. Lines used for measurements: JL001 x DR5revGFP, JL001 x DR5 VENUS, JL002 x DR5revGFP, and JL002 x DR5revVENUS (for PI4P); SB001 x DR5revGFP (for PI(4,5)P₂); SB037 x DR5revGFP and SB038 x DR5revGFP (for PS). See Table S1 for details of the lines.

This analysis revealed no detectable changes in PI4P or PI(4,5)P₂ levels from three hours before to two to three hours after FC division (Figure 3A-D). PS abundance also remained stable prior to FC activation but increased sharply afterwards within two hours (Figure 3E, F). These results indicate that PS undergoes a rapid post-activation increase, while PI(4,5)P₂ dynamics are slower and occur later during LRP development. Specifically, in Stage II LRPs, PI(4,5)P₂ levels decrease in the core compared to the flanks (Figure 1), although this change is not observed immediately after FC activation (Figure 3D). This suggests that spatial heterogeneity in PI(4,5)P₂ may play a more prominent role during LRP morphogenesis than during initiation.

### PI(4,5)P**₂** Is Required for Lateral Root Initiation

To investigate the functional role of PI(4,5)P₂ in LR development, we analysed LR formation in the *pip5k1 pip5k2* double mutant, which is deficient in PI(4,5)P₂ biosynthesis. PIP5K1 and PIP5K2 encode phosphatidylinositol 4-phosphate 5-kinases that are critical for auxin-mediated vascular patterning and primary root growth. Loss of both enzymes leads to a marked reduction in PI(4,5)P₂ levels (15, 33–36). However, their combined role in LR initiation and emergence had not been previously addressed. As expected, *pip5k1 pip5k2* seedlings exhibited severely reduced primary root growth (Figure 4A; Supplemental Figure S2), consistent with earlier reports. Unexpectedly, the combined density of LR and LRPs was significantly higher in the mutant than in the wild type (Figure 4B; Supplemental Figure S2). Closer examination revealed that this increase did not reflect enhanced LR initiation but instead resulted from a reduction in cell length. Quantification showed that fully elongated cortical cells in the mutant were approximately half the length of those in wild-type roots (Figure S2). To assess LR initiation independently of cell size, we used the Lateral Root Initiation (LRI) Index, which normalises the number of LR initiation events to a root segment containing 100 cortical cells (37). This analysis revealed that LR initiation was significantly reduced in the mutant, with the LRI reaching only ∼70% of the wild-type level (Figure 4B), indicating a requirement for PI(4,5)P₂ in efficient initiation of LRPs. We further evaluated LR emergence, defined as the proportion of LRPs that successfully develop into emerged LRs. In 6-day-old seedlings, emergence was significantly impaired in the *pip5k1 pip5k2* mutant: only 26% of LRPs progressed to emergence compared to 50% in wild-type (Figure S2). Together, these findings demonstrate that PI(4,5)P₂ is essential for promoting LRP initiation, underscoring a role for this lipid in the early phases of lateral root development.

**Figure 4.**
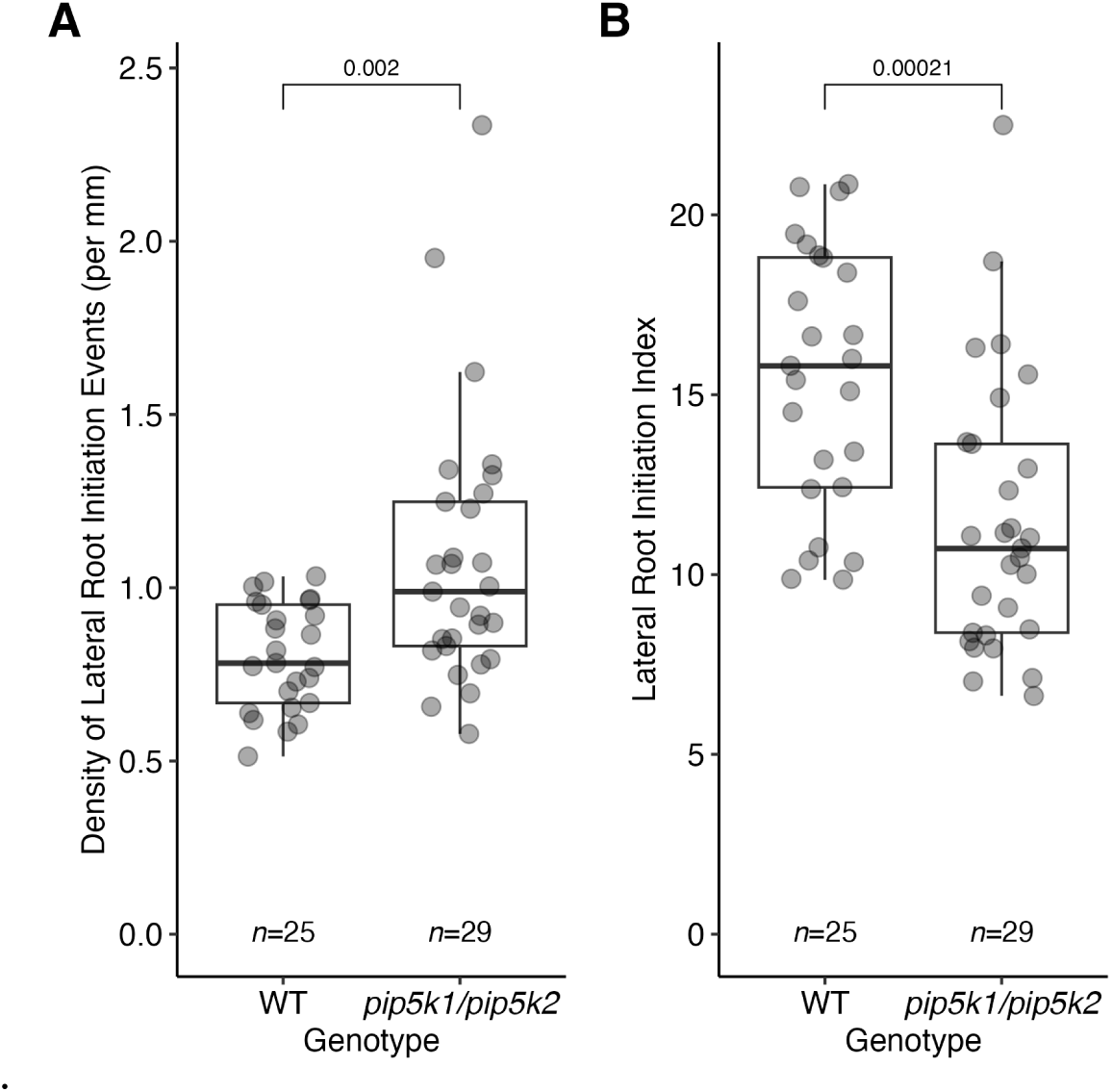
Lateral root initiation is significantly affected in *pip5k1pip5k2* double mutant. (A) Density of lateral root initiation events (number of initiation events per mm of root) in wild-type (WT) and *pip5k1pip5k2* double mutants. (B) Lateral Root Initiation Index, which normalises initiation events to a fixed number of cortical cells (100 cells per file), is significantly reduced in *pip5k1pip5k2* mutants compared to WT (*P* = 0.00021), indicating that true initiation frequency is lower in the mutant background. Each data point represents a single root; *n* indicates the number of roots analysed. *P*-values were calculated using a non-parametric Kruskal-Wallis one-way analysis of variance on ranks.

### Inducible Depletion of PI(4,5)P**₂** in Pericycle Cells Accelerates LRP Development Post Founder Cell Activation

To determine whether spatially targeted depletion of PI(4,5)P₂ can influence LR development, we employed the inducible iDePP system (38), which allows controlled depletion of PI(4,5)P₂ at the plasma membrane via a phosphatase. The system utilises the phosphatase domain of the *Drosophila melanogaster* OCRL protein (dOCRL), fused to a myristoylation/palmitoylation (MAP) membrane-targeting sequence and tagged with mCherry for visualisation (MAP-mCherry-dOCRL, in short iDePP, Figure 5A). A catalytically inactive version (iDePP_DEAD) served as a control. To assess the functional role of PI(4,5)P₂ specifically in pericycle-derived cells, we generated dexamethasone (DEX)-inducible iDePP lines under the control of two promoters with distinct spatiotemporal expression patterns. The *pXPP* promoter is active in xylem-pole pericycle (XPP) cells and in FCs but turns off when FCs are activated and form an LRP; at stage II, its activity is almost absent (Figure 5B). The *pGATA23* promoter is also active in XPP cells and FCs, but in contrast to *pXPP* it remains active until LRP stages III–IV (De Rybel et al., 2010; Figure 5C).

**Figure 5.**
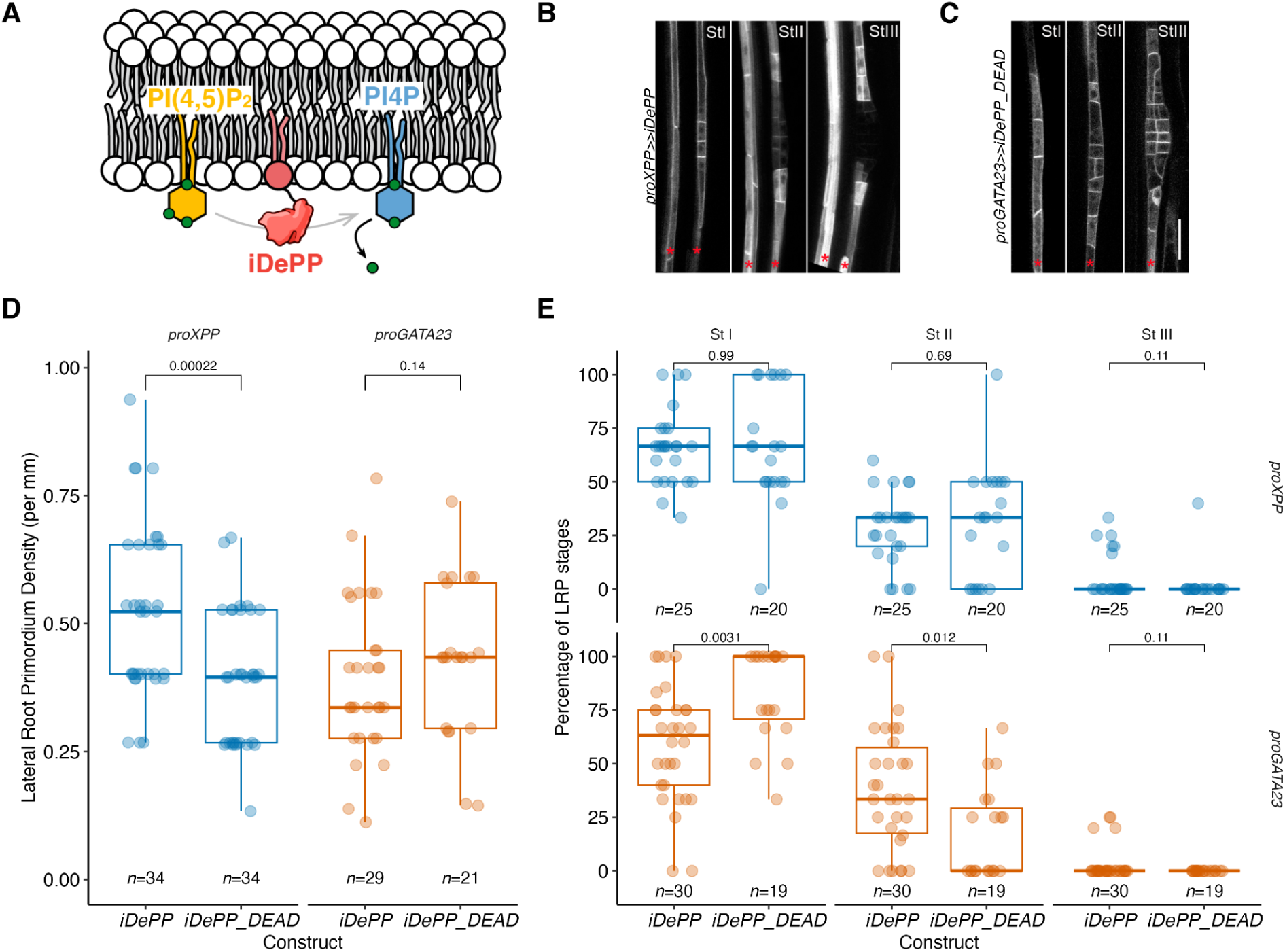
Inducible depletion of PI(4,5)P₂ stimulates LR formation. (A) Schematic representation of the iDePP system used for inducible depletion of PI(4,5)P₂ at the plasma membrane. The phosphatase domain of Drosophila OCRL, fused to mCherry, (iDePP, red) dephosphorylates PI(4,5)P₂ (yellow) to PI4P (blue). (B–C) Representative confocal images showing lateral root primordia (LRPs) at different developmental stages (St I–III) in seedlings inducibly expressing the iDePP construct under the *pXPP* (B) or *pGATA23* (C) promoters, after 7 and 6 h, respectively, of induction by 10 µM DEX. Asterisks mark XPP and LRPs. Scale bar = 40 µm, same for (B) and (C). (D) Quantification of LRP density (number of LRPs per mm of primary root) in iDePP and catalytically inactive control (iDePP_DEAD) lines driven by *pXPP* (blue) and *pGATA23* (orange) promoters induced by 10 µM DEX over a 24 h period. PI(4,5)P₂ depletion in XPP cells (*pXPP*) significantly increased LRP density (*P* = 0.00022), while no significant difference was observed in *pGATA23* lines (*P* = 0.14). (E) Distribution of LRP developmental stages (St I–III) in iDePP and iDePP_DEAD lines under *pXPP* (top) and *pGATA23* (bottom) promoters grown in the same conditions as in (D) . In *pGATA23* lines, PI(4,5)P₂ depletion led to a significant reduction in Stage I LRPs and a corresponding increase in Stage II and Stage III LRPs (*P* = 0.0031 and *P* = 0.012, respectively), indicating accelerated LRP progression. No significant stage differences were observed in *pXPP* lines. For all boxplots, individual data points are combined from two experiments, and *n* indicates the number of LRPs analysed. *P*-values from Kruskal-Wallis one-way analysis of variance on ranks are indicated.

To define optimal DEX treatment conditions, we first monitored the kinetics and localisation of iDePP. We found that 6–7 hours of DEX exposure was sufficient to activate the system, while prolonged treatment (beyond 31 h) led to mislocalization of the fusion protein into vacuoles—an effect observed in both iDePP and iDePP_DEAD lines (Figure S3). By 4 days post-induction, non-specific signal localisation was observed in all LRPs. To avoid confounding effects from ectopic localisation, we limited DEX treatment to a 24-hour period in all functional assays.

At 5 days after germination (dag), seedlings were transferred to medium containing 10 µM DEX for 2 hours and then rotated 90° to induce gravitropic stimulation. 24 hours after transfer, the newly formed root region was excised, cleared, and analysed. In *pGATA23* lines, root elongation during this time window was significantly greater in iDePP plants compared to iDePP_DEAD controls, suggesting that PI(4,5)P₂ depletion in founder-cell domains promotes root growth (Figure S3). In contrast, no difference in root elongation was observed in *pXPP* lines (Figure 5A, Figure S3). The number of LRPs formed during this 24-hour window was only slightly affected by PI(4,5)P₂ depletion, regardless the depletion took place in LRPs till stage III or earlier (Figure 5B). These data indicate that early PI(4,5)P₂ depletion in XPP cells (via *pXPP*) enhances LR initiation, while depletion throughout the pericycle and in early LRPs (via *pGATA23*) affects primary root growth but not LR initiation, possibly having a greater effect on LRP morphogenesis.

To assess whether PI(4,5)P₂ depletion influences the rate of LRP development, we staged the LRPs formed during the 24-hour period following induction. In *pGATA23* iDePP lines, a greater proportion of LRPs had progressed to Stage II or III compared to iDePP_DEAD controls (Figure 5D), indicating that depletion of PI(4,5)P₂ in founder-cell-derived cells and its derivatives accelerates primordium development. In contrast, PI(4,5)P₂ depletion in *pXPP* lines (taking place only in XPP and FCs) had no statistically significant impact on LRP stage progression (Figure 5E). Nevertheless, many more LRPs were found progressing to St III. Collectively, these results demonstrate that PI(4,5)P₂ acts as a negative regulator of LRP morphogenesis. Its targeted depletion enhances both LRP initiation (Figure 5D) and LRP development (Figure 5E), suggesting that PI(4,5)P₂ functions both in modulating the initiation of LRPs and in restraining their morphogenesis during early developmental stages.

## DISCUSSION

We investigated how the spatial and temporal dynamics of anionic phospholipids—particularly PI4P, PI(4,5)P₂, and phosphatidylserine (PS)—regulate LR formation in *Arabidopsis thaliana*. Using live-cell lipid biosensors, we found that PI4P is uniformly distributed across developing LRPs, while PI(4,5)P₂ exhibits a domain-specific depletion in the proliferative core. This pattern emerges during early primordium stages and is also reflected in XPP cells following LR initiation. PS, in contrast, accumulates at the plasma membrane following FC activation. Genetic analysis of *pip5k1pip5k2* mutants confirmed that PI(4,5)P₂ is required for LR initiation and emergence. Moreover, inducible depletion of PI(4,5)P₂ using the iDePP system revealed that localised reduction of this lipid in XPP, FCs, and early LRP stages accelerates LR initiation and LRP development. This study shows that dynamic regulation of anionic phospholipids, particularly PI(4,5)P₂ and PS, plays a central role in the spatial and temporal control of LR initiation and morphogenesis in *Arabidopsis thaliana*. Through biosensor imaging, mutant analysis, and inducible lipid manipulation, we provide evidence that changes in the abundance and localisation of these lipids are tightly coordinated during FC specification, activation, and primordium development.

Our spatial mapping of phosphoinositides during LR development showed that PI4P is evenly distributed across all LRP domains and remains stable before and after LR initiation. In contrast, PI(4,5)P₂ displayed domain-specific depletion: its levels decreased in the core of developing LRPs and in XPP cells following LR initiation, particularly at the outer and transverse plasma membrane domains. These findings suggest that PI(4,5)P₂ is specifically downregulated in highly proliferative cells, consistent with previous observations that LRP core cells exhibit a higher division rate compared to the slower-dividing flank cells (22, 23).

Interestingly, a similar spatial heterogeneity in phosphoinositide distribution has been reported in the shoot apical meristem (SAM), where PI(4,5)P₂ is depleted in the central zone and enriched at organ boundaries (39). While in the SAM, low PI(4,5)P₂ correlates with stem cell identity, in the root, reduced PI(4,5)P₂ is observed in actively dividing LRP core cells prior to quiescent centre establishment (40). Thus, rather than marking stemness, PI(4,5)P₂ depletion may instead reflect or facilitate cellular states associated with rapid proliferation and early morphogenetic transitions in LR development.

Our inducible depletion experiments using the iDePP system further support this view. When PI(4,5)P₂ was reduced only in XPP and FC, a greater LRP initiation was found, suggesting that PI(4,5)P₂ modulation is involved in LR initiation. When PI(4,5)P₂ was reduced in a broader domain (XPP, FC, and early LRP stages), we observed accelerated LRP progression and increased primordium stage advancement. This effect was not observed when PI(4,5)P₂ was depleted only in XPP cells and FCs, suggesting that PI(4,5)P₂ also functions as a negative regulator of morphogenesis. These results highlight the functional relevance of the endogenous PI(4,5)P₂ decrease observed in LRP core cells and imply that spatially restricted PI(4,5)P₂ depletion may help establish proliferative asymmetry within the developing primordium.

The role of PS adds another layer of complexity. PS, which contributes to plasma membrane electrostatics, showed a distinct response to LR initiation: its abundance increased specifically at the inner membrane of XPP cells and rose rapidly following FC activation. This timing and localisation suggest that PS may reinforce plasma membrane identity and polarity during early LRP development. Given that PS promotes auxin-dependent ROP6 signalling and stabilises ROP6 nanodomains (11), its enrichment during FC activation may help couple auxin response with cytoskeletal reorganisation and growth axis establishment. Our findings align with this model and provide evidence that PS acts in parallel with PI(4,5)P₂ to modulate membrane signalling domains crucial for LRP progression.

Genetic analysis of *pip5k1pip5k2* double mutants, which are deficient in PI(4,5)P₂ synthesis, revealed a contrasting phenotype to iDePP-mediated depletion. These mutants exhibited reduced LR initiation and impaired emergence, supporting the idea that while moderate, localised reduction of PI(4,5)P₂ is beneficial, global depletion disrupts auxin transport and polarity (33). This dichotomy emphasises the importance of precise spatial and temporal control of PI(4,5)P₂ levels. Notably, PI(4,5)P₂ is synthesised from PI4P by PIP5Ks, and its levels are likely balanced by local phosphatase activity. The observed effects of iDePP activation and the lack of a compensatory increase in PI(4,5)P₂ after FC activation suggest that a tightly regulated PI4P/PI(4,5)P₂ ratio may be essential for orchestrating key transitions in LRP morphogenesis.

Beyond cell division, PI(4,5)P₂ may also influence cell shape and growth axis establishment. In animal systems, PI(4,5)P₂-rich plasma membrane microdomains interact with microtubules to control membrane dynamics and cell motility (Golub and Caroni, 2006). A similar mechanism could operate in plants, where radial expansion of FCs accompanies early LRP formation. It is tempting to hypothesise that a local reduction in PI(4,5)P₂ relieves cytoskeletal constraints, enabling radial growth and changes in the cell polarity vector, processes critical for forming the dome-shaped structure of the primordium.

In summary, our work identifies PI(4,5)P₂ and PS as key regulators of lateral root development whose spatial dynamics are tightly linked to cell identity, polarity, and proliferative behaviour. PI4P appears to serve as a stable basal lipid pool, while PI(4,5)P₂ undergoes active regulation that likely integrates hormonal signalling, membrane electrostatics, and cytoskeletal organisation. These findings open new avenues to explore how lipid signalling contributes to developmental patterning and organogenesis in plants.

## MATERIALS AND METHODS

### Plant material and growth conditions

Plants of *Arabidopsis thaliana,* all in Col-0 background, were grown in vertically oriented Petri dishes under fluorescent illumination (100 µE m^-2^ s^-1^) in long-day conditions (16-h light/8-h dark) at 22°C. Seeds were surface sterilised (ethanol 70% and SDS 0.05%), stratified (4°C in the dark, > 24 h), and placed on 0.2x Murashige and Skoog (MS) medium adjusted to pH 5.7 (Duchefa) containing 0.8% Phyto Agar. For most experiments, Plants were analysed six to eight days after germination.

### Construction of vectors and plant transformation

All constructs were assembled using the GreenGate modular cloning system (41, 42). See Supplemental Table S2 for details. *Agrobacterium tumefaciens*-based plant transformation was done using the floral dip method (43).

### Histochemical analysis and microscopy

A Leica TCS SP5 laser scanning microscope equipped with HC PL APO 63x/1,20 W CORR and HC PL APO 40x/1,1 W CORR lenses was used. Fluorescent reporters were excited for GFP/YFP at 488/514 nm and for Scarlet/mCherry at 561/569 nm. Sequential and bidirectional scanning was used. The emission fluorescence for GFP/YFP and Scarlet/mCherry was collected in 493-583 nm and 570-680 nm windows. Roots were cleared using the method of Malamy and Benfey (1997) and mounted on slides as described (Dubrovsky et al., 2006). Identification of LRP was performed on cleared preparations under a Zeiss microscope, equipped with Nomarski optics. For analysis of cleared roots, a Plan-Apochromat 40x/0,95 objective was used.

### Lateral root quantification

Analysis of LR development was performed as described (44). In WT and *pip5k1pip5k2* mutants, the position of the most distal LRP was marked on a slide, and all initiation events were recorded. The combined density of LR and LRPs was evaluated per primary root portion from the root base to the location of the most distal LRP. The length of this root portion was measured with Fiji (45). LRI index was evaluated as described (Dubrovsky et al., 2009). Cortical cell length was measured with an ocular micrometre. When LRP protruded from the epidermis surface, it was considered as an emerged LR. The percentage of emerged LRs was evaluated as a ratio between the number of such roots to the total number of first-order LRs and LRPs per individual primary root. LRP developmental stages were recognised in accordance with Malamy and Benfey (1997).

### Time-lapse experiments

The following T3 lines were used: *pXPP::mScarlet:FAPP1:tHsp18.2:FastRed* (JL001) and *pGATA23::mScarlet:FAPP1:tHsp18.2:FastRed* (JL002) for PIP4; *pXPP::mScarlet:2XPH(PLC):FastGreen* (SB001) for PI(4,5)P₂; *pXPP::mScarlet:1XC2Lact:FastGreen* (SB037) and *pGATA23::mScarlet:1XC2Lact:FastGreen* (SB038) for PS. These lines were crossed with *DR5rev::GFP* or *pDR5::3xVENUS-N7*, (30, 31). F1 generation was used for time-lapse experiments, using in most cases 6-7 dag plants. In about half of the experiments, a 6 to 8 h gravitropic stimulation was applied that consisted of a 90 ° rotation of vertically oriented Petri dishes before the beginning of a time-lapse experiment. Plants were mounted in growth chambers in a 1-well Thermo Scientific™ Nunc™ Lab-Tek™ II Chamber Slide™ System. The seedling roots were covered with a 3 mm thick rectangular piece of 0.2×MS agar medium prepared as mentioned above, and the shoots were left in the air, free of agar. Experiments were of 11 to 16 h duration, and images were acquired for most experiments every 60 minutes (for some experiments, every 30 and 90 minutes). Time-lapse experiments were performed under a Leica TCS SP5 LSM equipped with a water immersion 63x 1.2 NA and 40x 0.8 NA objectives.

### Inducible depletion of PI(4,5)P**₂**

The following lines were used: *6xOP::iDePP DEAD-tHSP18.2M:KanR-pGATA23::GR-LhG4-tHSP18.2M* (JL004, used as a control in which no decrease in PI(4,5)P₂ is expected), *6xOP::iDePP-tHSP18.2M:KanR- pGATA23::GR-LhG4-tHSP18.2M* (JL006, used for inducible dephosphorylation of PI(4,5)P₂ in cells showing high pGATA23 promoter activity), *6xOP::iDePP DEAD-tHSP18.2M:KanR- pXPP::GR-LhG4-tHSP18.2M* (JL007, used as a control in which no decrease in PI(4,5)P₂ is expected) and *6xOP::iDePP-tHSP18.2M:KanR- pXPP::GR-LhG4-tHSP18.2M* (JL009, used for inducible dephosphorylation of PI(4,5)P₂ in cells showing high pXPP promoter activity). Five independent homozygous transformants of each line were tested for robust and tissue-specific induction of the iDePP system by monitoring mCherry fluorescence under a confocal microscope in 5-day-old seedlings germinated on 0.2x MS medium and transferred to a 10 µM DEX-containing medium for 12 h. These lines were then propagated to subsequent generations (up to T5). For phenotyping, seedlings were grown in 0.2x MS medium for five days after germination and then transferred to the same medium supplemented with 10 µM DEX. Root tip position was marked, and plants left to grow for 2 h in vertically oriented Petri dishes, after which they were 90 ° rotated for gravistimulation. Petri dishes were scanned, and root growth increment was measured using Fiji (45). Then the root portions formed during 24 h were excised with scissors under a binocular at a precise position of the mark, and the root portions were cleared by Malamy and Benfey (1997) method and analysed as described above.

### Image analysis

Signal intensities were measured using Fiji (45). Grey values were measured with an ROI across a plasma membrane parallel to the longitudinal cell wall of an XPP or FC. Analysing the changes in FCs, the fluorescent signal outside the root was measured as a background, and this value was subtracted from the measured signal intensity maximum value of the plasma membrane. The signal strength in the plasma membrane of FCs at the time when the first division was completed (time 0) was considered as a unit, and the measurements before and after the first division of a FC were expressed as a ratio to the signal strength at time 0.

### Statistical analysis

Plants were randomly assigned to groups and treatments during sowing. Sample size was not determined in advance; the largest possible number of plants per group was analysed. No blinding measure was installed. Statistical analyses and plotting were performed with R (v4.5.1). The methods and *P*-values are summarised in the figure legends. The code and data source files used for the generation of all the figures are available from https://github.com/Maizel-lab/Dubrovsky_et_al_Lipids.

## ACKNOWLEDGEMENTS

We are gratefully thankful for the sabbatical research support granted to J.G.D. by the Dirección General de Asuntos del Personal Académico (DGAPA) – Programa de Apoyos para la Superación del Personal Académico (PASPA) – Universidad Nacional Autónoma de México (UNAM) and for a research scholarship award from the German Academic Exchange Service (Deutscher Akademischer Austauschdienst, DAAD). This work was supported by the German Research Foundation (DFG) through the FOR2581 grants (MA5293/6-2) and the Consejo Nacional de Ciencia y Tecnología of Mexico (CONACYT) to BJRH (grant 740701). We thank N. Doktor, M.W. Garcia Jimenez, and J.M. Hurtado Ramírez for technical help and C. Hardtke for the *pip5k* mutant seeds.

## AUTHOR CONTRIBUTIONS

Conceptualisation: AM, JGD

Data acquisition and curation: JGD, EB, CG

Formal Analysis: AM, JGD

Funding Acquisition: AM, JGD

Methodology: JGD, JRH

Project Administration: AM

Resources: JL, SB

Supervision: AM, JGD

Figure creation: AM, JGD

Writing – Original Draft: AM, JGD

Writing – Review and Editing: AM, JGD

## DISCLOSURE AND COMPETING INTERESTS STATEMENT

The authors declare no competing interests.

## DATA AVAILABILITY

The data produced in this study are available from https://github.com/Maizel-lab/Dubrovsky_et_al_Lipids (quantifications, analysis scripts) and upon reasonable request (source images).

## Supplemental Material and Figures for

**Figure S1 (related to Figure 1).**
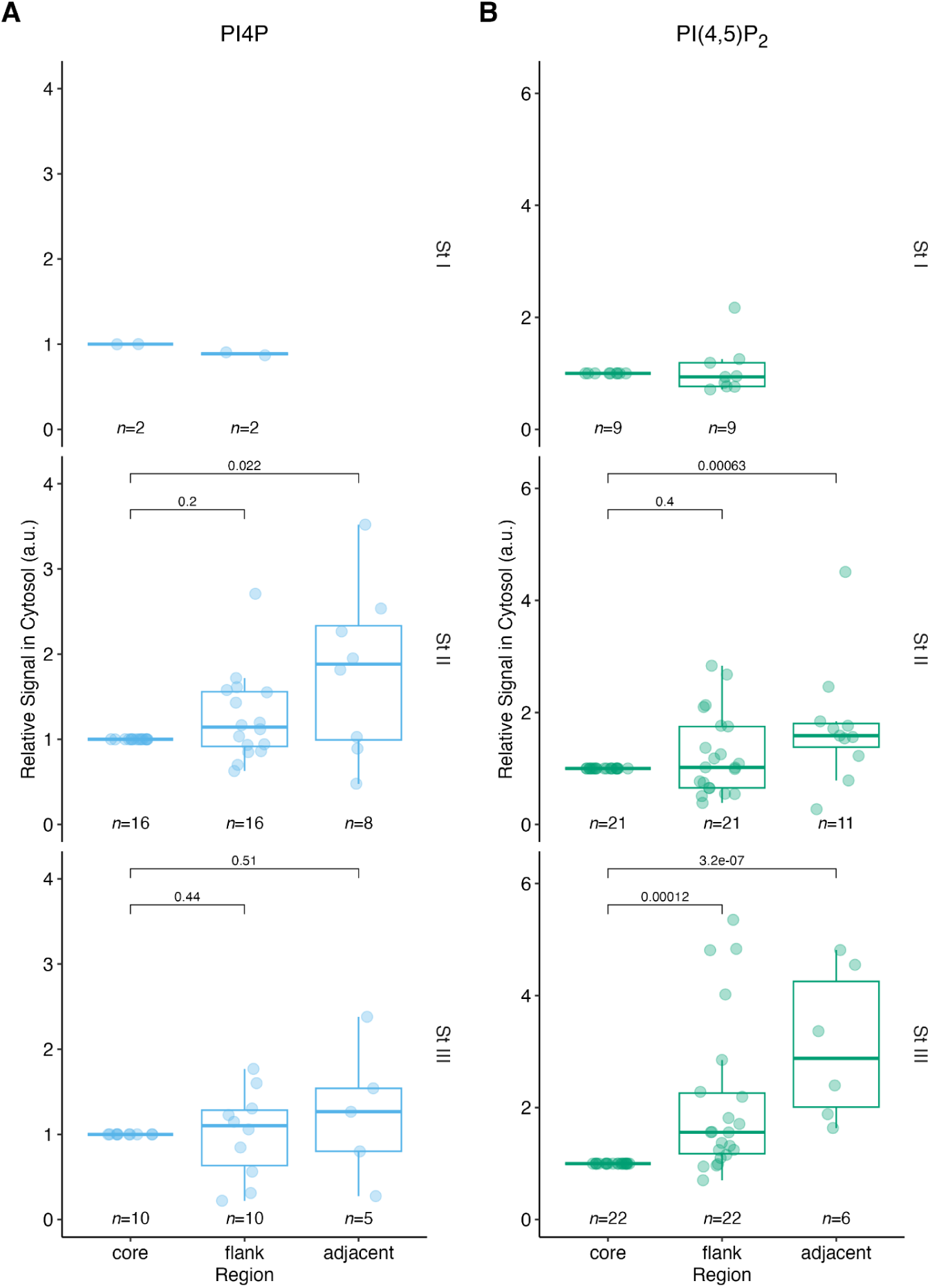
Cytoplasmic distribution of PI4P and PI(4,5)P₂ during LRP development. (A–B) Quantification of cytoplasmic signal intensity of PI4P (A) and PI(4,5)P₂ (B) biosensors in the core, flanking, and adjacent pericycle regions of lateral root primordia (LRPs) at developmental stages I to III. Signal intensity is expressed as relative fluorescence (a.u.) normalised to that in the LRP core. (A) PI4P cytoplasmic signal remains relatively uniform across compartments at the studied developmental stages, with no consistent trend of enrichment or depletion. (B) In contrast, PI(4,5)P₂ cytoplasmic signal is significantly reduced in the LRP core relative to that in adjacent cells at both, stage II (*P* = 0.00063) and stage III (*P* = 3.2e−07 and 0.00012); in stage III, the signal at core is also reduced compared to a flanking region (*P = 0.00012).* These changes reflect a pronounced and consistent depletion within the core region. Each dot represents a single cell; *n* indicates the number of cells analysed per group. Boxplots show medians and interquartile ranges. *P*-values were determined using a non-parametric Kruskal-Wallis one-way analysis of variance on ranks.

**Figure S2 (related to Figure 4).**
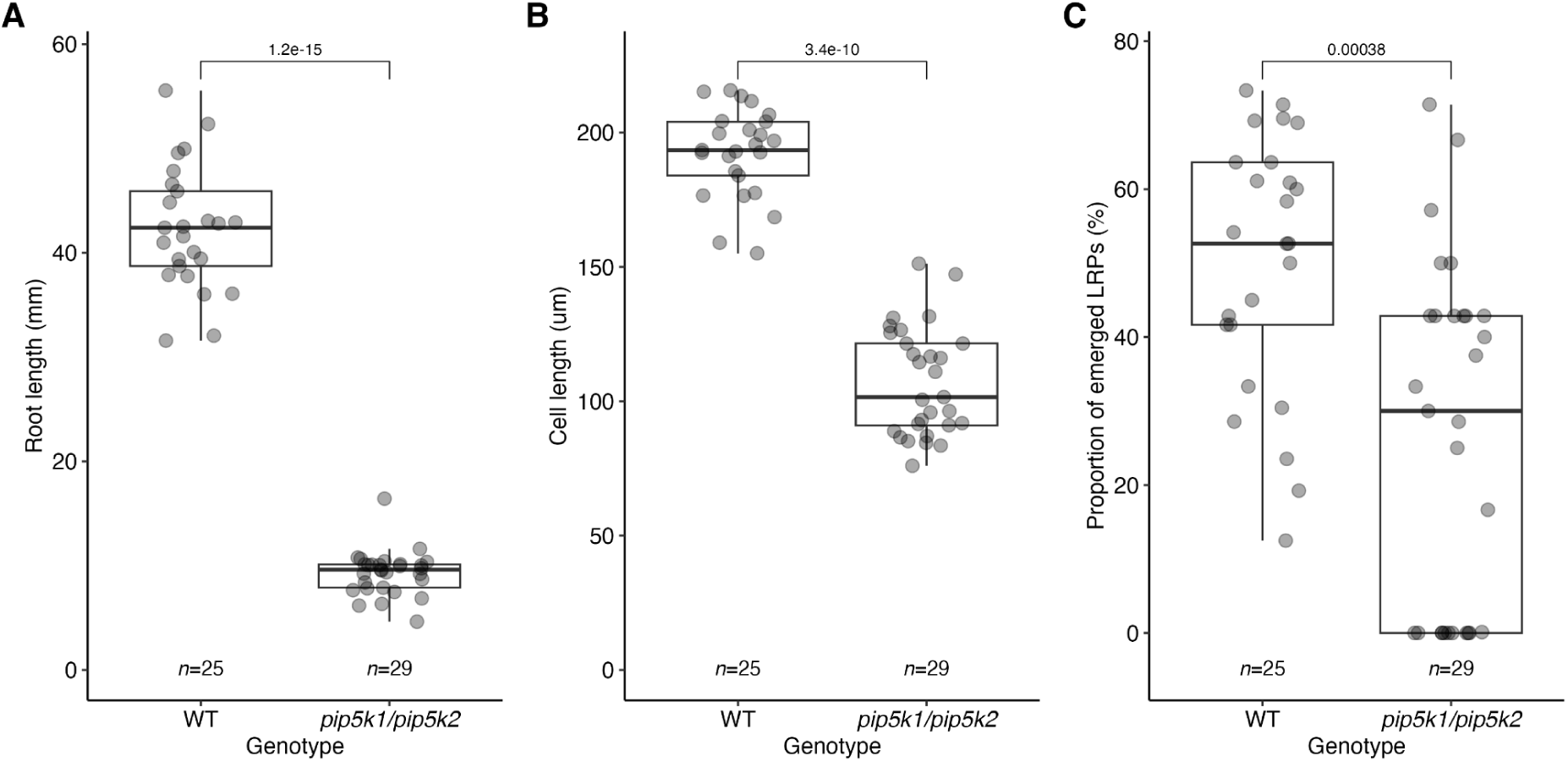
Quantitative analysis of lateral root development in the *pip5k1pip5k2* double mutant. Length of the primary root (A), of cortical cells (B) and proportion of emerged LR primordia (C) in WT and *pip5k1pip5k2* evaluated in 6-dag plants. Combined data of two independent experiments. *n* indicates the number of roots examined. Comparison was assessed by a non-parametric Kruskal-Wallis one-way analysis of variance on ranks. The *P* values are indicated.

**Figure S3 (related to Figure 5).**
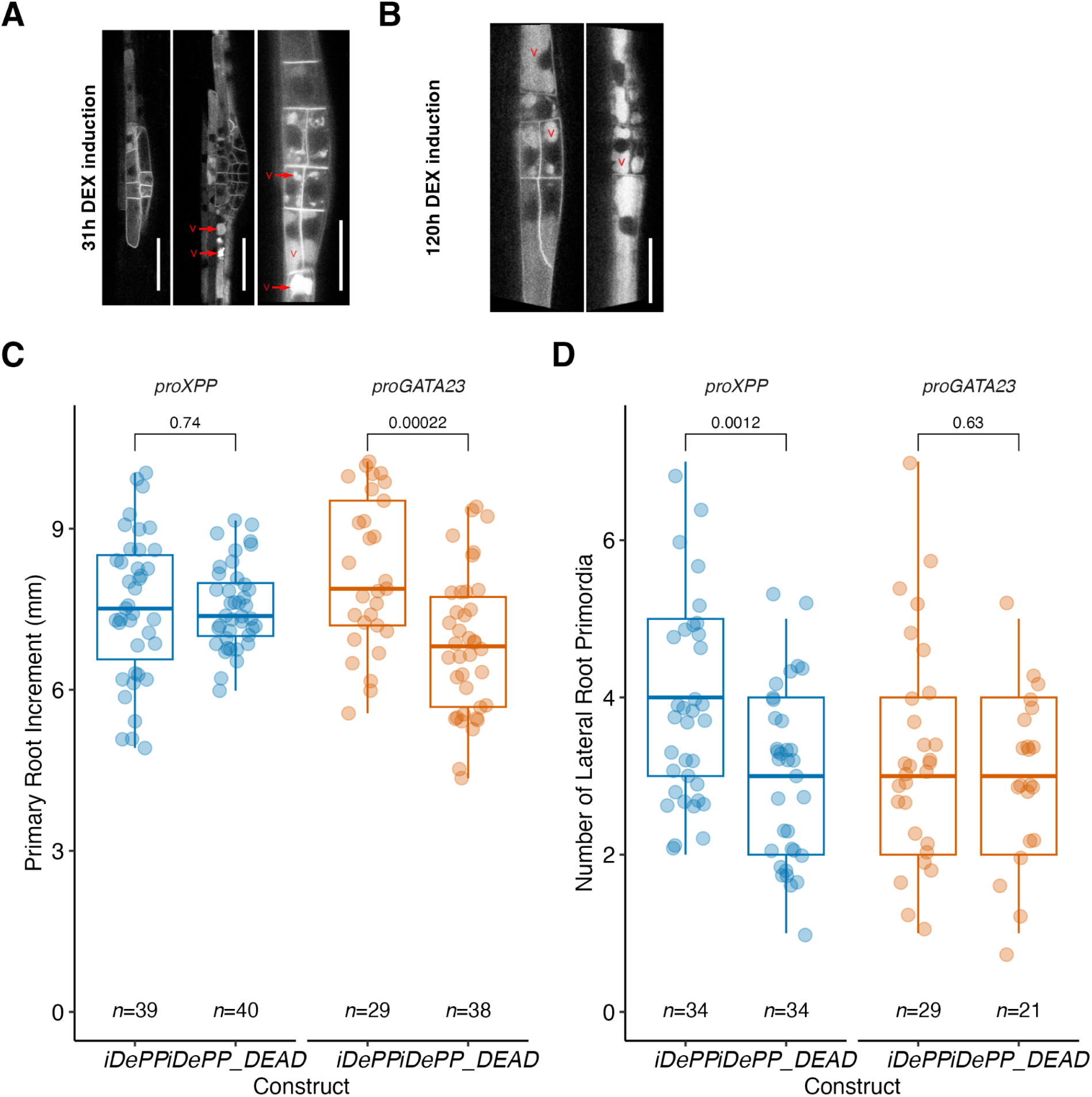
Domain-specific depletion of PI(4,5)P₂ differentially affects root elongation and LRP initiation. (A–B) Accumulation of the iDePP proteins in vacuoles upon sustained induction. Representative confocal images of root segments following DEX-induced activation of the iDePP system in *pGATA23* and *pXPP* lines. Roots imaged 31 hours (A) or 120 hours (B) after 10 µM DEX induction show vacuolar accumulation of the iDePP proteins (red arrows). Scale bars = 40 µm. (C) Quantification of primary root growth (increment in mm) 24 hours after DEX induction. In *pGATA23* lines (orange), iDePP expression significantly increased root growth compared to the catalytically inactive iDePP_DEAD control (*P* = 0.00022), while no significant difference was observed in *pXPP* lines (blue; *P* = 0.74). (D) Quantification of the number of LRPs formed during the 24-hour period following 10 µM DEX induction. PI(4,5)P₂ depletion in XPP cells (*pXPP*) significantly increased LRP number (*P* = 0.0012), whereas depletion in founder cells under *pGATA23* promoter did not (P = 0.63). For all boxplots, individual data points are combined from two experiments, and *n* indicates the number of plants analysed. *P*-values from Kruskal-Wallis one-way analysis of variance on ranks are shown.

**Table S1.** Plant lines used.

| Line Name | Construct | Background | Purpose | Figure | Reference |
| --- | --- | --- | --- | --- | --- |
| P7Y (sR782) | *pUBQ10::CITRINE-1xPH(OSBP)* | WT | PI4P biosensor | Figure 1 | Simon et al. 2014 |
| P24Y (sR785) | *pUBQ10::CITRINE-2xPH(PLC)* | WT | PI(4,5)P<sub>2</sub> biosensor | Figure 1 | Simon et al. 2014 |
| JL012 | *pRPS5A::mScarlet:2xFAPP/FastRed* | *pUBQ10:3xGFP:PIP1,4/*<br>*GATA23:H2B:3xmCherry* | PI4P biosensor | Figure 2 | This work |
| SB007 | *pXPP::mScarlet:2xPH(PLC)/FastGreen* | *pUBQ10:3xGFP:PIP1,4/*<br>*GATA23:H2B:3xmCherry* | PI(4,5)P<sub>2</sub> biosensor | Figure 2 | This work |
| SB008 | *pGATA23::mScarlet:2xPH(PLC)/FastGreen* | *pUBQ10:3xGFP:PIP1,4/*<br>*GATA23:H2B:3xmCherry* | PI(4,5)P<sub>2</sub> biosensor | Figure 2 | This work |
| SB009 | *pRPS5A::mScarlet:2xPH(PLC)/FastGreen* | *pUBQ10:3xGFP:PIP1,4/*<br>*GATA23:H2B:3xmCherry* | PI(4,5)P<sub>2</sub> biosensor | Figure 2 | This work |
| SB043 | *pXPP::mScarlet:1xC2Lact/FastGreen* | *pUBQ10:3xGFP:PIP1,4/*<br>*GATA23:H2B:3xmCherry* | PS biosensor | Figure 2 | This work |
| SB044 | *pGATA23::mScarlet:1xC2Lact/FastGreen* | *pUBQ10:3xGFP:PIP1,4/*<br>*GATA23:H2B:3xmCherry* | PS biosensor | Figure 2 | This work |
| SB045 | *pRPS5A::mScarlet:1xC2Lact/FastGreen* | *pUBQ10:3xGFP:PIP1,4/*<br>*GATA23:H2B:3xmCherry* | PS biosensor | Figure 2 | This work |
| SB037 | *pXPP::mScarlet:1xC2Lact/FastGreen* | WT | PS biosensor | Figure 3 | This work |
| SB038 | *pGATA23::mScarlet:1xC2Lact/FastGreen* | WT | PS biosensor | Figure 3 | This work |
| JL001 | *pXPP::mScarlet:2xFAPP/FastRed* | WT | PI4P biosensor | Figure 3 | This work |
| JL002 | *pGATA23::mScarlet:2xFAPP/FastRed* | WT | PI4P biosensor | Figure 3 | This work |
| SB001 | *pXPP::mScarlet:2xPH(PLC)/FastGreen* | WT | PI(4,5)P<sub>2</sub> biosensor | Figure 3 | This work |
| DR5revGFP (sR813) | *pDR5rev::GFP* | WT | Auxin signalling output reporter | Figure 3 | Benkova et al. 2003 |
| DR5revVENUS (sR242) | *pDR5rev::3XVENUS-N7* | WT | Auxin signalling output reporter | Figure 3 | Heisler et al. 2005 |
| *pip5k1 pip5k2* | - | - | phosphatidylinositol 4-phosphate 5-kinases mutant | Figure 4 | Ischebeck et al., 2013 |
| JL007 | *pXPP::LhG4>>6xOP::MAP-mCherry-dOCRL (iDePP)* | WT | Inducible PI(4,5)P<sub>2</sub> depletion | Figure 5 | This work |
| JL009 | *pXPP::LhG4>>6xOP::MAP-mCherry-dOCRL (iDePP_DEAD)* | WT | Control for iDePP experiments | Figure 5 | This work |
| JL006 | *pGATA23::LhG4>>6xOP::MAP-mCherry-dOCRL (iDePP)* | WT | Inducible PI(4,5)P<sub>2</sub> depletion | Figure 5 | This work |
| JL004 | *pGATA23::LhG4>>6xOP::MAP-mCherry-dOCRL (iDePP_DEAD)* | WT | Control for iDePP experiments | Figure 5 | This work |
| **Crosses** |  |  |  |  |  |
| SB001 x DR5revGFP | - | - | - | Figure 3 | This work |
| SB038 x DR5revGFP | - | - | - | Figure 3 | This work |
| SB037 x DR5revGFP | - | - | - | Figure 3 | This work |
| DR5revGFP x SB038 | - | - | - | Figure 3 | This work |
| JL001x DR5revGFP | - | - | - | Figure 3 | This work |
| JL001 x DR5revVENUS | - | - | - | Figure 3 | This work |
| JL002x DR5revGFP | - | - | - | Figure 3 | This work |
| JL002x DR5revVENUS | - | - | - | Figure 3 | This work |

**Table S2.** Transgenes generated.

| Construct | Plasmid | Assembly Type | Braided Level 1 plasmids | A | B | C | D | E | F | Destination |
| --- | --- | --- | --- | --- | --- | --- | --- | --- | --- | --- |
| *pXPP::mScarlet:2xFAPP/FastRed* | pJL001 | Level 1 | - | pAVB017 | pR0491 | pSB016 | pR0323 | pR0439 | pLB012 | pGGZ003 |
| *pGATA23::mScarlet:2xFAPP/FastRed* | pJL002 | Level 1 | - | pAVB004 | pR0491 | pSB016 | pR0323 | pR0439 | pLB012 | pGGZ003 |
| *pRPS5A::mScarlet:2xFAPP/FastRed* | pJL003 | Level 1 | - | pR0481 | pR0491 | pSB016 | pR0323 | pR0439 | pLB012 | pGGZ003 |
| *6xOP::MAP-mCherry-dOCRL (inactive)* | pJL007 | Level 1 | - | pR0368 | - | pJL004 | - | pR0439 | pR0303 | pMP51 |
| *6xOP::MAP-mCherry-dOCRL* | pJL009 | Level 1 | - | pR0368 | - | pJL006 | - | pR0439 | pR0303 | pMP51 |
| *proGATA23::LhG4* | pJL010 | Level 1 | - | pAVB004 | pR0283 | pR0572 | pR0323 | pR0439 | pMP59 | pMP52 |
| *pXPP::LhG4* | pJL011 | Level 1 | - | pAVB017 | pR0283 | pR0572 | pR0323 | pR0439 | pMP59 | pMP52 |
| *pXPP::mScarlet:2xPH(PLC)/FastGreen* | pSB003 | Level 1 | - | pAVB017 | pR0491 | pSB001 | pR0323 | pR0439 | pR0485 | pGGZ003 |
| *pGATA23::mScarlet:2xPH(PLC)/FastGreen* | pSB004 | Level 1 | - | pAVB004 | pR0491 | pSB001 | pR0323 | pR0439 | pR0485 | pGGZ003 |
| *pRPS5A::mScarlet:2xPH(PLC)/FastGreen* | pSB005 | Level 1 | - | pR0481 | pR0491 | pSB001 | pR0323 | pR0439 | pR0485 | pGGZ003 |
| *pXPP::mScarlet:1xC2Lact/FastGreen* | pSB010 | Level 1 | - | pAVB017 | pR0491 | pSB009 | pR0323 | pR0439 | pR0485 | pGGZ003 |
| *pGATA23::mScarlet:1xC2Lact/FastGreen* | pSB011 | Level 1 | - | pAVB004 | pR0491 | pSB009 | pR0323 | pR0439 | pR0485 | pGGZ003 |
| *pRPS5A::mScarlet:1xC2Lact/FastGreen* | pSB012 | Level 1 | - | pR0481 | pR0491 | pSB009 | pR0323 | pR0439 | pR0485 | pGGZ003 |
| *pGATA23::LhG4>>6xOP::MAP-mCherry-dOCRL (iDePP_DEAD)* | pJL012 | Level 2 (GreenBraid) | pJL007 pJL010 | - | - | - | - | - | - | pMP53 |
| *pGATA23::LhG4>>6xOP::MAP-mCherry-dOCRL (iDePP)* | pJL014 | Level 2 (GreenBraid) | pJL009 pJL010 | - | - | - | - | - | - | pMP53 |
| *pXPP::LhG4>>6xOP::MAP-mCherry-dOCRL (iDePP)* | pJL015 | Level 2 (GreenBraid) | pJL007 pJL011 | - | - | - | - | - | - | pMP53 |
| *pXPP::LhG4>>6xOP::MAP-mCherry-dOCRL (iDePP_DEAD)* | pJL017 | Level 2 (GreenBraid) | pJL009 pJL011 | - | - | - | - | - | - | pMP53 |

## Notes

https://doi.org/10.5281/zenodo.16778815

https://github.com/Maizel-lab/Dubrovsky_et_al_Lipids

### Competing Interest Statement

The authors have declared no competing interest.

### Summary of Updates

This revision includes the supplemental figures and table that were missing in the initial submission.

